# Weight cycling induces innate immune memory in adipose tissue macrophages

**DOI:** 10.1101/2022.07.02.498553

**Authors:** Heather L. Caslin, Matthew A. Cottam, Jacqueline M. Piñon, Likem Y. Boney, Alyssa H. Hasty

**Author notes:** Corresponding Author: Alyssa H. Hasty, PhD, 702 Light Hall, 23^rd^ Avenue South and Pierce, Nashville, TN 37232.

## Abstract

Weight loss improves obesity-associated diabetes risk. However, most individuals regain weight, which worsens the risk of developing diabetes and cardiovascular disease. We previously reported that male mice retain obesity-associated immunological changes even after weight loss, suggesting that immune cells may remember the state of obesity. Therefore, we hypothesized that cycles of weight gain and loss, otherwise known as weight cycling, can induce innate memory in adipose macrophages. We first treated bone marrow derived macrophages in a culture model of innate immune memory. Priming the cells with palmitic acid or adipose tissue conditioned media increased maximal glycolysis, oxidative phosphorylation, and increased LPS-induced TNFα and IL-6 production. While weight loss improved glucose tolerance, adipose macrophages retained elevated LPS-induced cytokine production. In a model of weight cycling, adipose macrophages had elevated metabolism and secreted higher levels of basal TNFα. Together, these data suggest that obesity and subsequent weight loss can prime adipose macrophages for enhanced inflammation upon weight regain. This innate immune memory response may contribute to worsened glucose tolerance following weight cycling.

## Introduction

Weight loss is effective for improving blood glucose, blood pressure, and blood lipids^1–3^. However, weight loss is hard to accomplish and even harder to maintain, and most individuals regain weight within a few years^4–7^. We refer to the repeated process of gaining and losing weight as weight cycling, which can occur throughout one’s lifetime. Unfortunately, weight cycling further increases the risk for developing type 2 diabetes, cardiovascular disease, and hypertension^8–11^, and has a stronger relationship with allcause mortality than even stable long-term obesity^12^. However, the mechanism by which weight cycling worsens disease risk remains unknown.

Obesity is a state of chronic, low-grade, systemic inflammation, and adipose immune cells contribute to obesity-associated disease. Upon weight gain, inflammatory cells expand and infiltrate into the adipose tissue and release inflammatory cytokines such as IL-1β, TNFα, and IL-6^13–18^. These cytokines directly promote adipocyte lipolysis and impair insulin signaling, contributing to the development of diabetes^19^. We previously published our work demonstrating that male C57BL/6J which undergo weight cycling have worsened glucose tolerance compared with their obese counterparts, and this is correlated with an increase in memory T cells in their adipose tissue^20^. We have also reported that while weight loss normalizes glucose tolerance, it does not restore obesity-associated immunological changes such as T cell exhaustion or macrophage lipid handling^21^. These data suggest that adipose immune cells may “remember” the state of obesity.

The induction of immune memory has long been known to be a key feature of T cells and B cells. However, recent studies have also revealed a memory response in innate cells, in which stimuli prime innate immune cells to augment subsequent activation to a second stimulus. This response, coined “innate immune memory” or “trained innate immunity”, was initially observed with β-glucan and the *Mycobacterium tuberculosis* vaccine (BCG), but has also been seen with cytokines, hormones, and oxidized low density lipoprotein^22–28^. Functionally, innate immune memory is associated with elevated glycolytic metabolism and inflammatory function. Mechanistically, this memory response is controlled by epigenetic modifications that hold open regions of chromatin at glycolytic and inflammatory genes^28^. We hypothesized that weight cycling induces innate immune memory in adipose tissue macrophages and could contribute to worsened disease risk.

In the present study, we show that previous exposure to palmitic acid or adipose conditioned media *in vitro* and weight loss *in vivo* increase macrophage metabolism and inflammatory cytokine production. This immunological memory may influence weight maintenance or the metabolic consequences of further weight gain.

## Methods

### Animals

Male and female C57BL/6J mice were either purchased from Jackson Labs or bred within our group. Additionally, TLR4 KO mice on a C57BL/6 background were provided by Dr. Brad Grueter at Vanderbilt Universtiy^29^. All procedures were approved and carried out with approval from and in compliance with the Vanderbilt University Institutional Animal Care and Use Committee. Vanderbilt University is accredited by the Association for Assessment and Accreditation of Laboratory Animal Care International.

### Weight cycling

For weight loss and weight cycling experiments, male C57BL/6J mice were purchased from Jackson Labs (#000664) at 7 weeks of age. At 8 or 9 weeks of age, mice were placed on 9-week cycles of high fat diet (60% fat, Research Diets #D12492, 5.21 kcal/g food) or low fat diet (10% fat, Research Diets #D12450B, 3.82 kcal/g) for a total of 27 weeks as published^20,21^ and as visualized in **Figure 1A**. Access to food and water was provided *ad libitum.* Body weight and food intake were recorded weekly.

**Figure 1:**
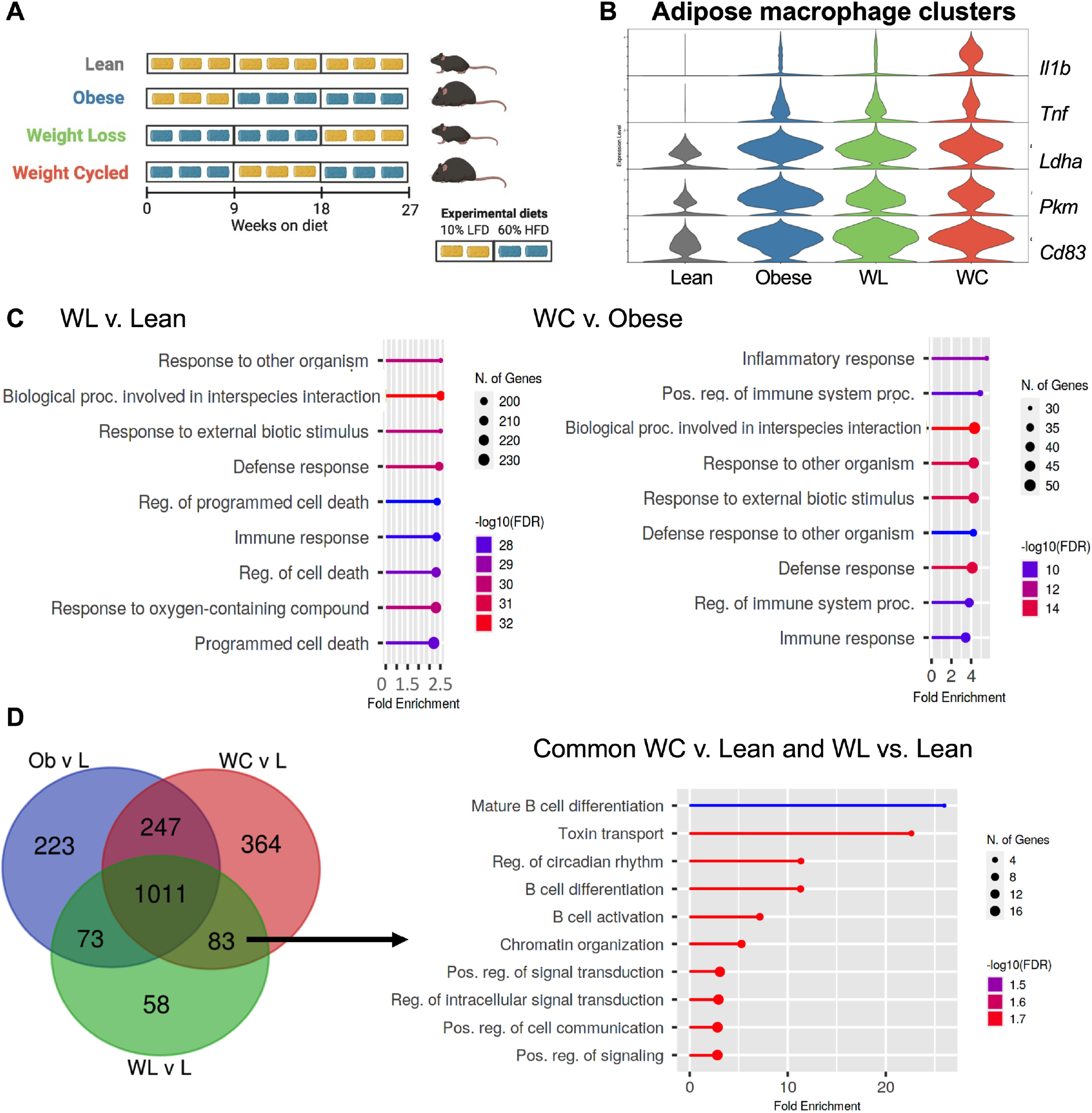
Transcriptional changes in adipose macrophages from weight loss and weight cycled mice suggest innate immune memory. **(A)** Weight cycling schematic. **(B)** Violin plot of inflammatory (*Tnf, Il1b, Cd83*) and metabolic genes (*Ldha, Pkm*) from lean, obese, weight loss (WL), and weight cycled (WC) macrophage clusters. **(C)** Pathway analysis of differentially expressed genes from WC vs. obese or WL vs. lean macrophage clusters. **(D)** Venn diagram of differentially expressed genes of obese, WL, and WC adipose macrophage clusters compared with lean and pathway analysis of the genes differentially expressed in both WL and WC groups.

### Single cell analysis

To assess adipose macrophage populations for innate immune memory by single cell-sequencing, we used a previously published single cell sequencing dataset from our laboratory (GEO accession number GSE182233)^21^. Differential expression was performed in R version 4.1 with Seurat V4^30^ using Wilcoxon Ranked Sum tests. GO Biological Processes pathway analysis was conducted on differential expression with ShinyGO 0.76^31^ using FDR cutoff of 0.05 (last date of access 6/15/2022) and overlapping gene expression was determined and graphed using http://bioinformatics.psb.ugent.be/webtools/Venn/.

### Bone marrow derived macrophages and adipose tissue conditioned media

For innate immune memory culture models, male and female mice on chow diets were euthanized between 8-15 weeks of age and bone marrow was extracted from the femurs. Bone marrow-derived macrophages (BMDM) were differentiated over 5-7 days in DMEM (Gibco, 11960044 or 11885092), 10% FBS, 1% Penicillin/streptomycin, 10 mM HEPES, and 10% L929 conditioned media (or 10 ng/mL M-CSF; Shenandoah Biotech # 200-08) in T75 flasks.

For adipose tissue conditioned media, male mice fed 60% high fat diet (Research Diets #D12492) for 5-9 weeks were euthanized and epididymal adipose fat pads were collected and minced. Small pieces of adipose tissue were cultured in DMEM with FBS, Pen/strep, and HEPES as above for 48 hours at ~50 mg/mL. Media was filtered through 50 μm filters to remove adipocytes, centrifuged to remove red blood cells and immune cells, and stored at −20°C for further experiments.

### In vitro innate memory model

For *in vitro* innate immune memory experiments, we adopted a cell culture model by Kleinnijenhuis, Quintin, and colleagues^23,24^. Briefly, BMDM were treated for 24 hours with 0.4 mM palmitic acid (MP Biomedicals #57-10-3) or 100% adipose tissue conditioned media. Palmitic acid was suspended in DMSO or conjugated to BSA. Media was washed out for 6 days and cells were stripped, re-plated at 375 cells/μL, and activated with 100 ng/mL LPS (Sigma #L4391), 1μg/mL lipotechoic acid (InvivoGen #tlrl-pstla), 1 μg/mL poly(I:C) (Tocris #4287), or 5 μg/mL beta-glucan (Millipore #346210) for 24 hours.Seahorse metabolic analysis was conducted at the end of the washout phase and cell culture media was collected at the end of 24-hour activation.

### Seahorse metabolic analysis

To measure cell metabolism, a Seahorse XFe96 (Agilent) was used to measure the extracellular acidification rate (ECAR) and oxygen consumption rate (OCR) as surrogates for glycolysis and oxidative phosphorylation, respectively. Cells were plated at 50-75,000/well in 200 μL of standard growth media for 2-24 hours and then switched to minimal DMEM containing 2 mM L-glutamine for the assay. A modified version of the Mito Stress Test (Agilent #103015-100) was then performed. Briefly, glucose (final concentration 10 mM) was injected to the wells to measure glucose-stimulated glycolysis. Pyruvate (final concentration 1 mM) was then injected with oligomycin (final concentration 1.5 μM) to begin the standard Mito Stress Test. Last, FCCP (final concentration 2 μM) and rotenone/antimycin A (final concentration 0.75 μM) were injected. To normalize for any differences in cell number due to treatment, cells were lysed in RIPA buffer and we performed a Pierce BCA assay (Thermofisher #23225) according to the manufacturers’ protocol. OCR and ECAR were normalized to μg protein.

### ELISA

To measure cytokine production, cell culture media was collected for ELISA. IL-6 and TNF murine ELISA kits were purchased from Biolegend (#431304, #430904). Assays were performed in duplicate when possible, according to the manufacturers’ protocols. To normalize for any differences in cell number due to treatment, cells were lysed in RIPA buffer and we performed a Pierce BCA assay (Thermofisher #23225) according to the manufacturers’ protocol. Cytokine concentrations were normalized relative to control samples from each experiment.

### Body composition

To assess body composition, mouse body fat and fat free mass (FFM) were measured by nuclear magnetic resonance whole body composition analysis (Bruker Minispec).

### Glucose tolerance testing

To assess glucose tolerance, mice were fasted for 5 hours and briefly anesthetized with isoflurane for a tail snip to access blood supply. After a 1-hour recovery, basal blood glucose levels were measured using a hand-held glucometer (Bayer Contour Next EZ meter). After an intraperitoneal injection of 1.5 g dextrose/kg lean mass, blood glucose was sampled at 15, 30, 45, 60, 90, and 120 minutes.

### Tissue immune cell isolation

To isolate adipose macrophages, the adipose stromal vascular fraction was collected as previously described^32^. Briefly, mice were euthanized by isoflurane overdose and cervical dislocation and perfused with 20 ml PBS through the left ventricle. Epididymal (unless otherwise stated) or subcutaneous adipose depots were collected, minced, and digested in 6 ml of 2-mg/mL type II collagenase (Sigma # C6885 or Worthington #LS004177) for 30 min at 37°C. Digested tissue was then vortexed, filtered through 100 μm filters with cold PBS, lysed with ACK buffer, and filtered through a 35 μm filter. The stromal vascular fraction was plated at 200,000 cells/well for adherence for 2-4 hours for Seahorse and ELISA experiments or stained for flow cytometry.

To isolate blood monocytes, whole blood was lysed in water and 500,000 cells/ well were plated for 2 hours for adherence and used for cytokine production.

To isolate peritoneal macrophages, peritoneal lavage was performed. Cold PBS (5 mL) with 5mM EDTA was injected into the abdominal cavity, the abdomen was massaged for 2 min, and lavage fluid was collected from a small incision by transfer pipette. 200,000 cells/well were plated for adherence for 2 hours for Seahorse metabolic analysis and cytokine production.

To isolate liver macrophages, livers were extracted, minced, and digested as above. The liver suspensions were then plunged through 100 μm filters with cold PBS and lysed with ACK buffer. The cell pellet was resuspended in 33% Percoll and overlayed on top of 66% Percoll. The Percoll gradient was centrifuged at 600 × g for 15 min with the break set to zero. The two middle layers of the Percoll gradient were collected in HBSS and centrifuged at 500 × g for 5 min. 200,000 cells/well were plated for 2 hours for adherence and used for Seahorse metabolic analysis and cytokine production.

### Flow Cytometry

To assess cell frequency and intracellular cytokine production, stromal vascular cells were prepared for flow cytometry. FcBlock was added for 10 min prior to fluorescent antibody staining for surface proteins for 30 min at 4°C using the following antibodies: BV510 anti-mouse CD45 (Biolegend 103137), FITC anti-mouse CD11b (eBioscience 11-0112-86), PerCP-Cy5.5 anti-mouse CD64 (Biolegend 139308), PE-Cy7 anti-mouse CD9 (Biolegend 124815), APC-Cy7 anti-mouse F480 (Biolegend 123118). eFlour 450 fixable viability dye (Invitrogen/eBioscience 50-112-8817 was used to determine viability). For intracellular cytokines, cells were fixed and permeabilized according to the Thermofisher two-step protocol: for intracellular (cytoplasmic) proteins (Foxp3/transcription factor staining buffer set, eBioscience 00-5523-00). Cells were then stained with fluorescent antibodies for intracellular proteins for 30 min at 4°C using the following antibodies: PE anti-mouse TNF (Biolegend 506306) and APC anti-mouse IL-6 (Biolegend 503810). Data was acquired on a MACSQuant10 (Miltenyi) and analyzed on FlowJo.

### Statistical analyses

Statistical analyses were performed using GraphPad Prism. Student’s *t*-tests were run for comparisons between two groups, one-way analysis of variances (ANOVA)s were used for comparisons between more than two groups, and two-way ANOVAs were conducted for >2 groups over >2 timepoints. For significant main effects, *post-hoc* pairwise comparisons using Tukey or Sidak corrections were used to determine statistical differences. All data are presented as mean ± standard error (SEM). A *p*-value (or adjusted *p*-value) of <0.05 was used to determine significance.

## Results

### Transcriptional induction of innate immune memory in adipose macrophages from weight loss and weight cycled mice

To determine if weight cycling can induce innate immune memory, we first utilized a recently published single cell dataset from our lab^21^. Briefly, mice had been placed on low fat or high fat diets for 27 weeks total (**Figure 1A**). At the end of 27 weeks, obese and weight cycled mice were matched for body weight, and obese, weight loss, and weight cycled mice were matched for total time on high fat diet. Single-cell sequencing was completed on CD45^+^ adipose tissue immune cells and analyses across many different cell populations were reported^21^. To extend upon those findings, genes from the Trained Immunity DataBase^33^ were plotted across our diet groups in the macrophage clusters (**Figure 1B**). We performed differential expression analysis and found inflammatory genes (*Il1b* and *Tnf*), glycolytic genes (*Pkm* and *Ldha*), and the activation marker *Cd83* were more highly expressed in adipose macrophages from weight loss compared to lean mice (adjusted p<0.05 by differential expression, **Supplemental Table 1**). Additionally, *Il1b* and *Cd83* were more highly expressed in the adipose macrophages from weight cycled vs. obese mice (adjusted p<0.05 by differential expression, **Supplemental Table 2**). Next, we performed pathway analysis on all significant differentially expressed genes to determine which pathways were differently regulated (**Figure 1C**). In weight loss compared with lean as well as weight cycled compared with obese comparisons, pathways related to immune regulation, inflammation, defense, and responses to biotic, external stimuli, and interspecies interaction were changed. These pathways have also been shown to change following the induction of innate immune memory by β-glucan or BCG^34,35^. To further determine the pathways changed in both weight loss and weight cycled groups, we used a Venn diagram to compare the differentially expressed genes in all groups when compared to the lean group (**Figure 1D**). Of the 83 genes that were significantly different in both the weight loss and weight cycled groups, common pathways related to chromatin organization, signal transduction, and cell communication. These cellular processes are all generally considered to be characteristic of innate immune memory. Interestingly, we also found pathways associated with differentiation and activation in B cells and toxin transport. Together, these data supported our hypothesis that weight loss and weight cycling may induce innate immune memory in adipose macrophages.

### Transcriptional induction of innate immune memory in monocytes and dendritic cells

Using the dataset from Figure 1, we also completed the same analyses on the monocyte and dendritic cell (DC) clusters (**Figure S1A & B**). *Il1b* was significantly increased in classical monocytes from weight cycled vs. obese mice (**Supplemental 3 & 4),** and both *Il1b* and *Tnf* were more highly expressed in the cDC2 and monocyte derived DC populations from weight cycled vs. obese (adjusted *p*-value <0.05, **Supplemental Table 5 & 6**). In the classical monocytes, pathways related to immune regulation, defense, responses to biotic or external stimuli were changed similar to adipose macrophages in Figure 1B and to the response following β-glucan or BCG^34,35^ (**Figure S1C**). Notably, cytokine response genes and metabolic processes were also changed. In the cDC2 and monocyte derived DC clusters, many of the pathways with the greatest change were related to translation, protein biosynthesis/metabolism, and cytokine response. In the clusters from weight cycled vs. obese mice, differences were observed in pathways related to innate receptor signaling and inflammation. These data further support the hypothesis that weight cycling may induce innate immune memory in innate myeloid cells of the adipose tissue.

### Palmitic acid induces innate immune memory responses in bone marrow derived macrophages

We next adopted a previously published cell culture model of innate immune memory with a 24-hour prime, 6-day washout, and 24-hour secondary activation (**Figure 2A**)^23,24^. Palmitic acid was chosen as our initial stimulus because it is the most common saturated fatty acid in the human body, is elevated in individuals with obesity due to lipolysis, and it activates TLR4, a receptor that can promote innate immune memory^36^. Additionally, palmitate has been shown to elicit an obese-like adipose macrophage phenotype in bone marrow derived macrophages (BMDM) and priming in other models^37–39^. Following 24-hour activation and a 6-day washout, palmitic acid priming increased maximal extracellular acidification rate (ECAR), a proxy for glycolysis, and appeared to increase maximal oxygen consumption rate (OCR), a proxy for oxidative phosphorylation, although this measure was not significant (**Figure 2B&C**). Moreover, following 24-hour LPS activation, palmitic acid priming increased LPS-induced TNFa and IL-6 secretion when compared to the control (**Figure 2D**). These results were observed when palmitic acid was suspended in either DMSO (shown) or BSA (data not shown). Cytokine production was also generally increased with palmitic acid priming in response to additional secondary stimuli such as lipoteichoic acid, a TLR2 agonist (**Figure 2E**), as well as β-glucan, a dectin-1 agonist (**Figure 2F**), although there were cytokine specific effects. We did not detect an increase in response to poly(I:C), a TLR3 agonist, suggesting that the priming effect influences a variety of, but not all, stimuli (**Figure 2E**). Additionally, there was no significant difference between palmitic acid primed or control BMDM activated with β-glucan in TLR4 KO BMDM (**Figure 2F**), suggesting that this response is dependent on palmitic acid activation of TLR4. As expected, there was no LPS activation in the TLR4 KO (data not shown). Together, these data suggest that palmitic acid increases metabolic potential and cytokine production to a secondary stimuli, which is consistent with other innate immune memory stimuli^23,24^.

**Figure 2:**
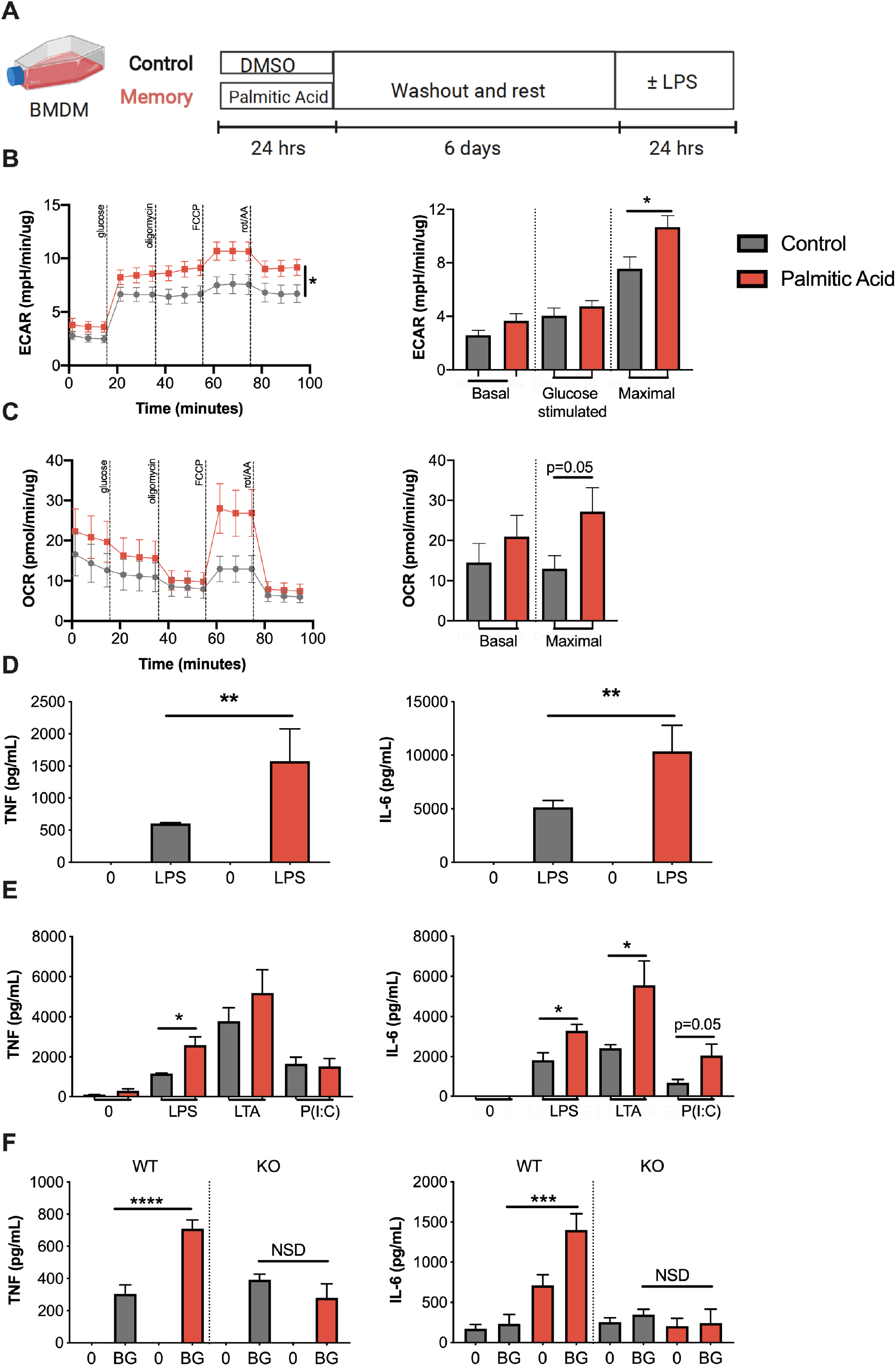
Palmitic acid increases maximal cell metabolism and inflammation in macrophages in culture. **(A)** Schematic of innate immune memory *in vitro* models using bone marrow derived macrophages and.4 mM palmitic acid or control (DMSO). **(B)** Extracellular acidification rate (ECAR) and **(C)** oxygen consumption rate (OCR) during modified mitochondrial stress test by Seahorse metabolic analyzer after 6 day rest. **(D)** 100 ng/mL LPS-induced TNF and IL-6 production by ELISA and normalized by protein concentration. **(E)** 100 ng/mL LPS-, 5 ug/mL LTA-or 5ug/mL P(I:C)-induced TNF and IL-6 production by ELISA and normalized by protein concentration. **(F)** 100 ng/mL LPS-induced TNF and IL-6 production by ELISA in WT or TLR4-KO mice and normalized by protein concentration. Data are means ± SEM of 3-7 populations, representative of 2-3 independent experiments. \*\**p* <.01, *** *p* <0.001, \*\*\*\**p* <.0001, NSD= no significant difference.

Lev Becker’s group has previously published that 24-hour treatment of BMDMs with palmitate induces metabolically activated macrophages (MMe) with elevated gene and protein expression of Plin2, Abca1, and Cd36 that is consistent with adipose macrophages from obese mice^37,38^. To confirm that our model of enhanced LPS-responsiveness is distinct from MMe activation, we compared 24-hour treatment with priming as above. Mme activation did not increase LPS-induced IL-6 or TNFa (**Figure S2**), suggesting that our innate immune memory model, and potentially weight cycling, is immunologically distinct from models of simple obesity.

### Adipose tissue conditioned media increases cell metabolism and inflammation in bone marrow derived macrophages

Innate immune memory has been shown in response to pattern recognition receptors like TLRs, but also in response to aldosterone, catecholamines, glucose, and cytokines^27,40–43^. Thus, we extended our above model to a more physiological model for weight cycling by priming cells with adipose tissue conditioned media (**Figure 3A**). Priming BMDM with adipose tissue conditioned media significantly increased maximal glycolysis and oxidative phosphorylation (**Figure 3B & C**). Priming with adipose tissue conditioned media also increased LPS-induced TNFα and IL-6 secretion (**Figure 3D**). These data further confirm that repeated exposure to an obese adipose tissue environment increases metabolic potential and cytokine production, which is again consistent with other innate immune memory stimuli.

**Figure 3:**
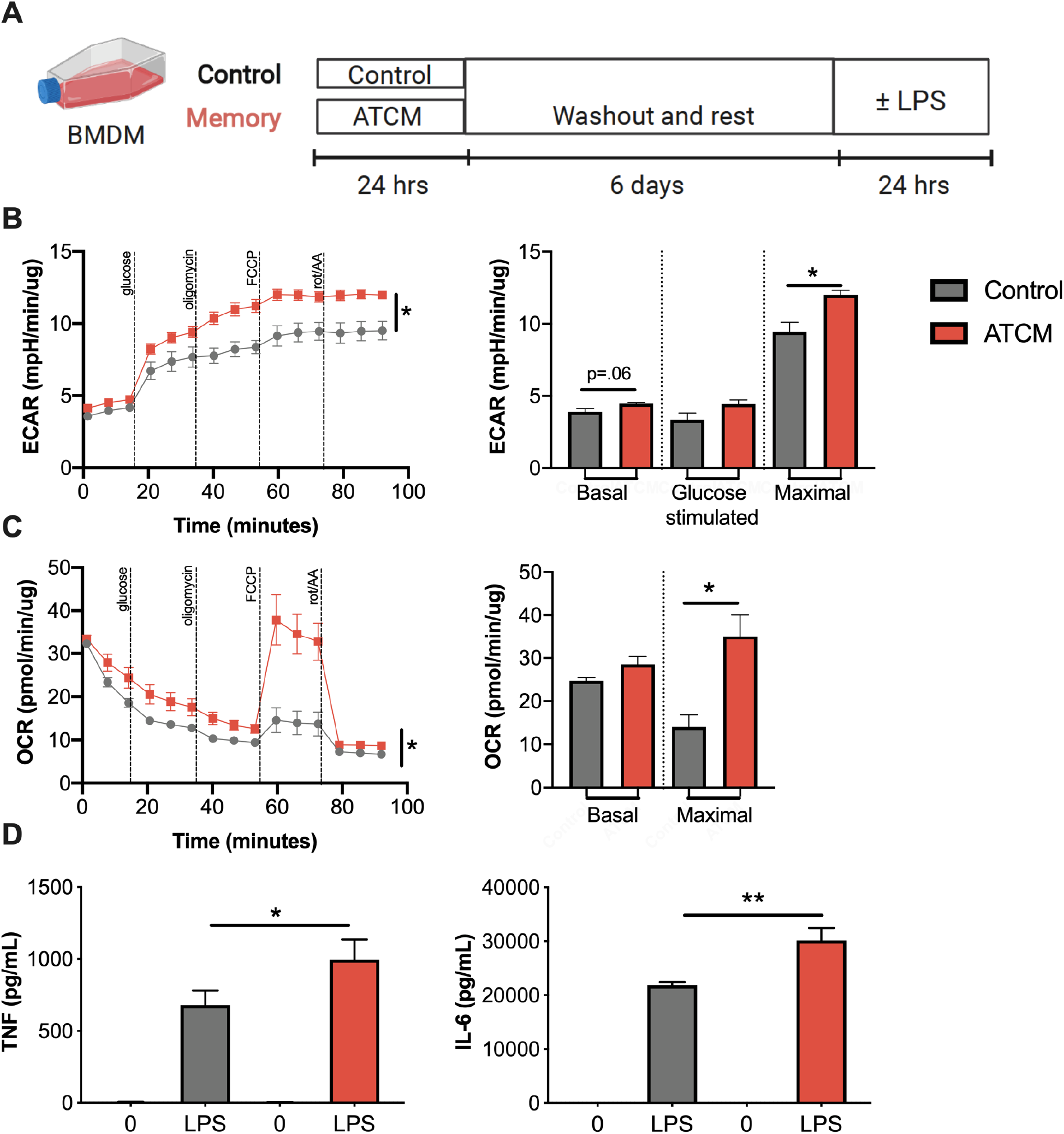
Adipose conditioned media increases cell metabolism and inflammation in macrophages and DCs in culture. **(A)** Schematic of innate immune memory *in vitro* models using bone marrow derived macrophages and adipose tissue conditioned media (ATCM) or control media. **(B)** Extracellular acidification rate (ECAR) and **(C)** oxygen consumption rate (OCR) during modified mitochondrial stress test by Seahorse metabolic analyzer. **(D)** 100 ng/mL LPS-induced TNF and IL-6 production by ELISA and normalized by protein concentration. Data are means ± SEM of 3 populations, representative of 2-3 independent experiments. \**p* <0.05, \*\**p* <.01.

### Weight loss mimics innate immune memory in adipose macrophages

We next tested if weight loss could replicate the effects observed when priming cells with palmitate and ATCM above. Mice were placed on low fat diet or high fat diet as in Figure 4A. After 9 weeks of high fat diet, we observed the predicted increase in body mass, fat mass, and lean mass in obese mice (**Figure 4B & C**). After switching to a low fat diet for 5 weeks, our weight loss mice had similar body weight and body composition to lean mice (**Figure 4B & C**). During an intraperitoneal glucose tolerance test, obese mice had higher blood glucose and slower glucose disposal at all timepoints compared to lean mice, however glucose tolerance was not significantly different in weight loss mice compared to lean mice (**Figure 4D**). Obesity increased glycolysis and oxidative phosphorylation in adipose macrophages as expected^44,45^ (**Figure 4E & F**). Maximal glycolysis appeared to remain elevated in adipose macrophages from weight loss mice, although this did not reach statistical significance. Importantly, adipose macrophages from weight loss mice increased LPS-induced TNFα and IL-6 production (**Figure 4G**). Blood monocytes displayed a similar response, with weight loss increasing LPS-induced IL-6 production (**Figure 4H**). Interestingly, these results were tissue specific, as peritoneal macrophages and liver macrophages had similar metabolism and cytokine production across groups (**Figure S3**). Moreover, in an extended model of weight loss (18 weeks), adipose macrophages still trended towards increased metabolic and inflammatory parameters, although only LPS-induced TNFα production was statistically elevated compared to the lean group (**Figure S4**). Together, these data suggest that while weight loss improves glucose tolerance, it increases inflammatory cytokine production in previously obese adipose macrophages. Moreover, these data suggest that weight loss derived signals may act as priming stimuli to elicit changes in adipose macrophage metabolism and cytokine production that are consistent with innate immune memory.

**Figure 4:**
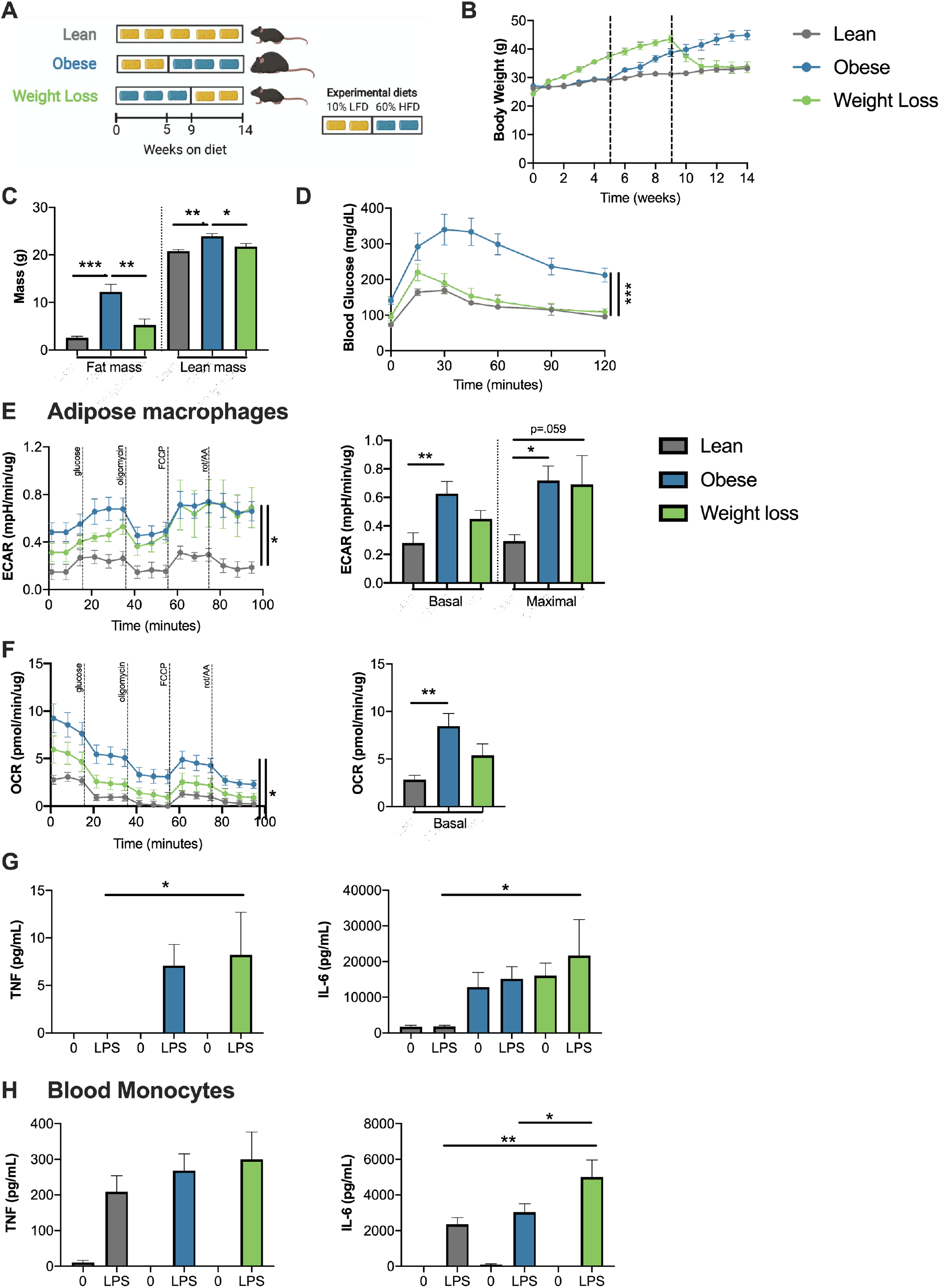
Weight loss doesn’t normalize the inflammatory response in adipose macrophages. **(A)** Weight loss *in vivo* schematic. **(B)** Body mass over time measured weekly with diet switch indicated by dashed lines. **(C)** Lean and fat mass measured by nuclear magnetic resonance. **(D)** Blood glucose during an intraperitoneal glucose tolerance test (1.5 g dextrose/kg lean mass) at 14 weeks. **(E)** Extracellular acidification rate (ECAR) and **(F)** oxygen consumption rate (OCR) of epididymal adipose macrophages selected by 2 hr adherence during modified mitochondrial stress test by Seahorse metabolic analyzer. **(G)** 100 ng/mL LPS-induced TNF and IL-6 production over 24 hours by ELISA in epididymal adipose macrophages selected by 2 hr adherence and normalized by protein concentration**. (H)** LPS-induced TNF and IL-6 production over 24 hours by ELISA in monocytes selected by 2 hr adherence and normalized by protein concentration. Data are means ± SEM of 6-8 mice, representative of 2 independent experiments. \**p* <0.05, \*\**p* <.01, *** *p* <0.001.

### Weight cycling further increases inflammation in adipose macrophages

To determine if weight regain provides a sufficient secondary activation signal for innate memory in vivo, we utilized our previously published model of weight cycling as shown in Figure 1A^20,21^. As previously reported, obese vs. weight cycled mice and lean vs. weight loss mice had comparable body weight, food intake and body composition at the end of 27 weeks (**Figure 5A – C**). As we previously published, weight cycling also worsened glucose tolerance, to a greater degree than obesity alone, following an intraperitoneal glucose tolerance test as measured by area under the curve (**Figure 5D**). While there were no differences in epididymal adipose macrophage proportions in obese vs. weight cycled mice (**Figure S5A & B**), weight cycling increased basal glycolysis and oxidative phosphorylation in adipose macrophages compared to lean macrophages (**Figure 5E & F**). Maximal metabolism was significantly elevated compared to lean macrophages in both the obese and weight cycled groups, with weight cycled trending higher. Basal TNFα production from epididymal adipose macrophages was also elevated in the weight cycled condition compared with lean macrophages (**Figure 5G**). We also observed an increase in intracellular TNFα in weight cycled epididymal adipose macrophages by flow cytometry (**Figure S5C**). Moreover, subcutaneous adipose macrophages and liver macrophages have increased TNFa production following weight cycling (**Figure S6A & B**). Together, these data suggest that weight loss induces a form of innate immune memory that enhances cell metabolism and TNFα production following weight regain.

**Figure 5:**
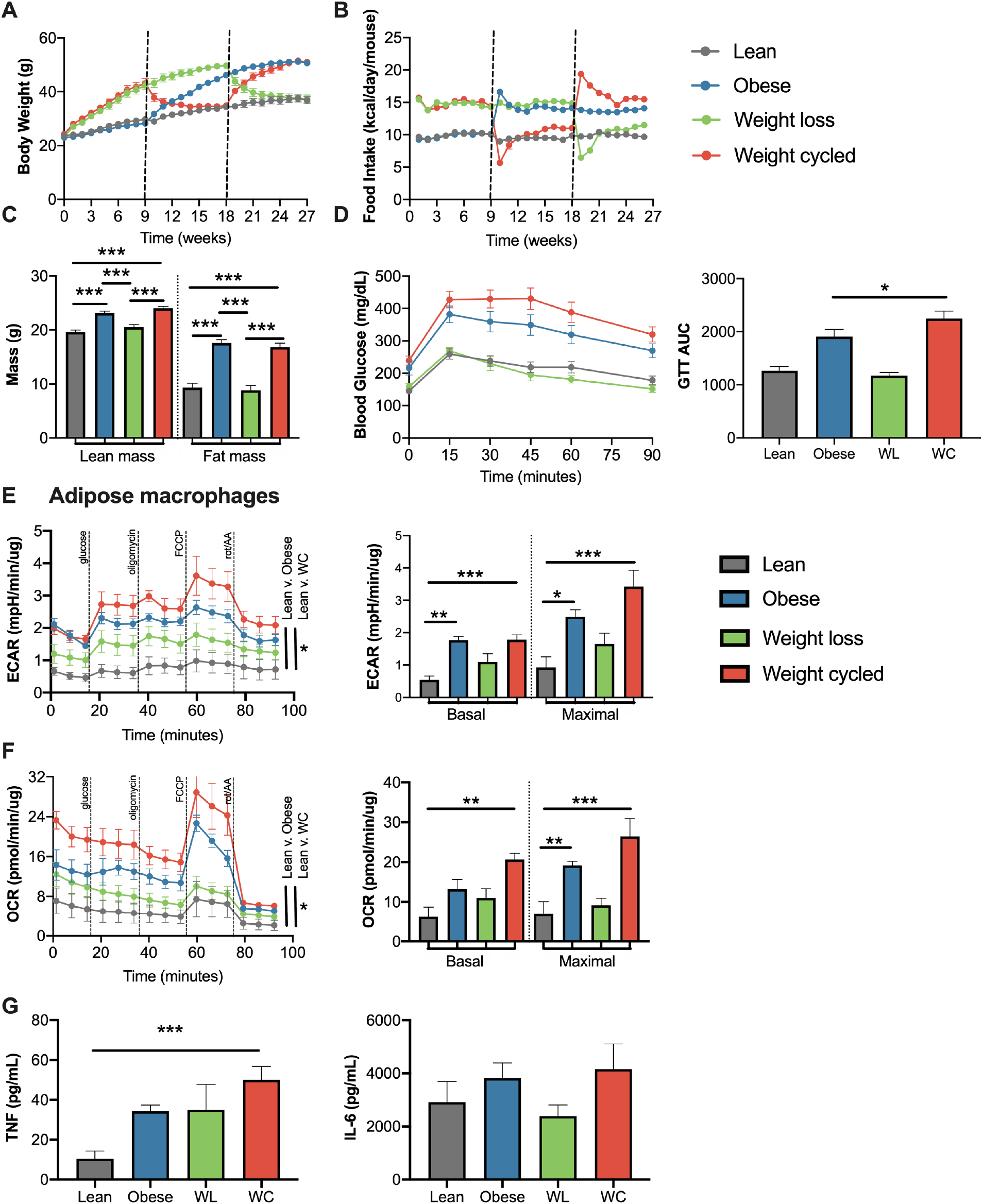
Weight cycling augments cell metabolism and basal cytokine production from adipose macrophages. Lean, obese, weight loss (WL) and weight cycled (WC) mice were generated over 27 weeks as in Figure 1A. **(A)** Body mass over time measured weekly with diet switch indicated by dashed lines. **(B)** Food intake over time measured weekly. **(C)** Lean and fat mass measured by nuclear magnetic resonance. **(D)** Blood glucose during an intraperitoneal glucose tolerance test (1.5 g dextrose/kg lean mass) at 27 weeks and area under the curve. **(E)** Extracellular acidification rate (ECAR) and **(F)** oxygen consumption rate (OCR) of epididymal adipose macrophages selected by 2 hr adherence during modified mitochondrial stress test by Seahorse metabolic analyzer. **(G)** TNF and IL-6 production over 24 hours by ELISA in adipose macrophages selected by 2 hr adherence and normalized by protein concentration. Data are means ± SEM of 10-12 mice for A-D, 5-14 mice for E-G, representative of 2 independent experiments. *\*p* <0.05, \*\**p* <.01, *** *p* <0.001.

### Discussion

In the current study, we demonstrate that previous exposure to palmitic acid or adipose conditioned media in culture, as well as in weight loss and weight cycled mice, exacerbates the inflammatory response of adipose macrophages. We consider this a metabolic form of innate immune memory, similar to that previously shown in the context of diabetes and atherosclerosis^46–48^. To our knowledge, this is the first study to link changes in adiposity with innate immune memory.

While weight cycling worsens diabetes disease risk beyond stable long-term obesity; the mechanisms that link weight cycling and disease risk are not known. Using single-cell sequencing, we previously showed that after weight loss, adipose immune cells retain obesity-associated immunological changes like T cell exhaustion and macrophage lipid handing^21^. Additionally, work by Lumeng and colleagues has shown that adipose macrophages from mice that have lost weight have greater inflammatory gene expression by microarray analysis^49^. In complementary work, oxidized LDL and Western diet increase LPS-responsiveness in monocytes and this is retained after 4 weeks of control diet^48^. Our current work supports these studies by showing that weight loss does not reverse enhanced LPS-induced cytokine production by adipose macrophages. Importantly, both Lumeng’s and our work show that these changes persist with long-term weight loss (18-24 weeks), suggesting these changes are long lasting and not simply in response to acute adipose tissue lipolysis or remodeling.

We propose this heightened inflammatory response is mediated by a metabolic form of innate immune memory. Initial reports of innate immune memory following exposure to β-glucan or BCG showed elevated glycolytic metabolism and inflammatory function^23,50^. We demonstrate similar changes in both cell culture and animal models, observing an increase in glycolysis and LPS-induced cytokine production. Priming with palmitic acid increases the response to many innate stimuli, including TLR2, TLR4, and dectin-1 agonists. Napier and colleagues have shown that other models of palmitate treatment can augment macrophage inflammation in culture and induce an innate memory-like effect^39^. Moreover, her group showed that dietary palmitate can augment the inflammatory response to endotoxemia and improve the clearance of *Candida albicans in vivo*. Together, our data emphasize the notion that palmitate induced-innate immune memory is not specific to one specific antigen, but primes macrophages for a more general heightened inflammatory response^28^. In our study, we also observed an increase in maximal oxidative metabolism, which has been shown following monophosphoryl lipid-A priming, which is a TLR4 agonist like palmitic acid^51,52^. Future studies should explore if palmitic acid or adipose tissue conditioned media induce any epigenetic modifications, which could be responsible for the sustained inflammatory phenotype following stimulus removal^28^.

Importantly, we also show that weight loss induces this metabolic form of innate immune memory *in vivo,* which may exacerbate diabetes risk upon weight regain. We showed that adipose macrophages produce more TNFα after weight regain, which could be one plausible link. Macrophage cytokine production can directly promote adipocyte lipolysis and impair insulin signaling in obesity^19^, which suggests a local mechanism by which weight cycling may impair glucose tolerance. However, recent experiments within our group suggest weight cycling does not worsen peripheral insulin resistance compared to obese animals, but rather impairs pancreatic insulin secretion (unpublished data, manuscript in revision). It is therefore plausible that secreted factors could reach other organs like the pancreas to promote dysfunction. Even more provocative, immune cells could possibly migrate from adipose tissue to other tissues. Alternatively, weight loss mobilizes lipid release from the adipose tissue^53^, and it is possible that metabolic macrophage memory development could occur in tissues like the pancreas. While we did not assess pancreatic macrophages in this study, we show that liver macrophages have increased TNFα production following weight cycling. A third possibility is that innate immune memory may be initiated in the bone marrow prior to recruitment upon weight regain, and recruited cells may also infiltrate the pancreas. Future work will directly interrogate the role of enhanced macrophage inflammation as a mechanistic link between weight cycling and worsened glucose tolerance.

There are a few noteworthy limitations to this study. First, we only used male mice for weight cycling studies. We expect to see similar inflammatory changes in female mice because we used both male and female BMDM for cell culture experiments; however, our current weight cycling model does not induce significantly worsened glucose tolerance in females^21^. Young female mice have very little weight gain and adipose expansion upon an initial exposure to high fat diet, and thus, the magnitude of initial priming may be important. A new model using aging, different diet lengths, or gonadectomy should be explored. Second, it is not known if metabolic memory is observed in human weight cycling or the extent to which humans would need to lose and gain weight. Finally, our model relies on a switch from high fat to low fat diets. We expect other models of weight loss with lipid mobilization to prime adipose macrophages; however, we have not assessed macrophage activation following other models of weight loss like calorie restriction or bariatric surgery. Future experiments should work to further identify metabolic memory in female mice, humans, and following different types of weight loss, including weight loss following gastric bypass, drug therapy, exercise, or other diets.

Taken together, our data add to the growing body of literature which shows that while weight loss restores systemic glucose tolerance, it does not restore the adipose immune landscape. We also identify palmitic acid, adipose conditioned media, and weight loss as stimuli for innate immune memory development. This is physiologically important because weight regain worsens risk for diabetes and other diseases, and adipose macrophage metabolic memory may be one mechanistic link for this association. Future studies should investigate the causal role of metabolic memory and find potential therapeutic targets to alleviate weight cycling-accelerated disease.

## Acknowledgements

This project was funded by a Veterans Affairs Merit Award 5I01BX002195 and an AHA Innovation Award (19IPLOI4760376) to AHH. HLC is funded by an AHA Postdoctoral Fellowship (20POST35120547), MAC is funded by an NIH F31 Predoctoral Fellowship (1F31DK123881), and both were previously supported by the Molecular Endocrinology Training Program (T32 DK07563). Most metabolic analyses were performed with the Seahorse Extracellular Flux Analyzer housed and managed within the Vanderbilt High-Throughput Screening Core Facility, an institutionally supported core, and was funded by NIH Shared Instrumentation Grant 1S10OD018015. Additional metabolic analyses were performed using Jeff Rathmell’s instruments. We would also like to thank Hasty Lab members Nathan Winn, Magdalene Ameka, Katie Volk, Monica Bhanot, Alec Rodriguez, and Marnie Gruen for their help with tissue collection and animal care during our studies. Figure schematics were created with Biorender.com.

## Contributions

HLC conceptualized the study, obtained and analyzed the data, and drafted the manuscript. MAC donated tissue from a few mouse cohorts and assisted with study design, sequencing data collection, and data interpretation. JP and LB assisted with culture model troubleshooting, data collection, and sequencing analysis. AHH assisted with study design, provided funding, and is the guarantor for this work. All authors contributed to manuscript revisions.

## References

1. Vidal, J. Updated review on the benefits of weight loss. Int J Obes Relat Metab Disord 26 Suppl 4, S25–28 (2002).

2. Ryan, D. H. & Yockey, S. R. Weight Loss and Improvement in Comorbidity: Differences at 5%, 10%, 15%, and Over. Current obesity reports 6, 187 (2017).

3. de Leiva, A. What are the benefits of moderate weight loss? Exp Clin Endocrinol Diabetes 106 Suppl 2, 10–13 (1998).

4. Wing, R. R. & Phelan, S. Long-term weight loss maintenance. Am J Clin Nutr 82, 222S–225S (2005).

5. Fildes, A. et al. Probability of an Obese Person Attaining Normal Body Weight: Cohort Study Using Electronic Health Records. Am J Public Health 105, e54–59 (2015).

6. Crawford, D., Jeffery, R. W. & French, S. A. Can anyone successfully control their weight? Findings of a three year community-based study of men and women. Int. J. Obes. Relat. Metab. Disord. 24, 1107–1110 (2000).

7. Stunkard, A. The Results of Treatment for Obesity: A Review of the Literature and Report of a Series. A.M.A. Archives of Internal Medicine 103, 79 (1959).

8. Bangalore, S. et al. Body-Weight Fluctuations and Outcomes in Coronary Disease. New England Journal of Medicine 376, 1332–1340 (2017).

9. Delahanty, L. M. et al. Effects of Weight Loss, Weight Cycling, and Weight Loss Maintenance on Diabetes Incidence and Change in Cardiometabolic Traits in the Diabetes Prevention Program. Diabetes Care 37, 2738–2745 (2014).

10. Montani, J.-P., Schutz, Y. & Dulloo, A. G. Dieting and weight cycling as risk factors for cardiometabolic diseases: who is really at risk? Obes Rev 16 Suppl 1, 7–18 (2015).

11. Rhee, E.-J. et al. Increased risk of diabetes development in individuals with weight cycling over 4 years: The Kangbuk Samsung Health study. Diabetes Research and Clinical Practice 139, 230–238 (2018).

12. Rzehak, P. et al. Weight change, weight cycling and mortality in the ERFORT Male Cohort Study. Eur. J. Epidemiol. 22, 665–673 (2007).

13. Weisberg, S. P. et al. Obesity is associated with macrophage accumulation in adipose tissue. Journal of Clinical Investigation 112, 1796–1808 (2003).

14. Russo, L. & Lumeng, C. N. Properties and functions of adipose tissue macrophages in obesity. Immunology 155, 407–417 (2018).

15. Ferrante, A. W. The immune cells in adipose tissue. Diabetes, Obesity and Metabolism 15, 34–38 (2013).

16. Jang, J. E. et al. Nitric Oxide Produced by Macrophages Inhibits Adipocyte Differentiation and Promotes Profibrogenic Responses in Preadipocytes to Induce Adipose Tissue Fibrosis. Diabetes 65, 2516–2528 (2016).

17. Thomas, D. & Apovian, C. Macrophage functions in lean and obese adipose tissue. Metabolism 72, 120–143 (2017).

18. Bing, C. Is interleukin-1β a culprit in macrophage-adipocyte crosstalk in obesity? Adipocyte 4, 149 (2015).

19. Saltiel, A. R. & Olefsky, J. M. Inflammatory mechanisms linking obesity and metabolic disease. J Clin Invest 127, 1–4 (2017).

20. Anderson, E. K., Gutierrez, D. A., Kennedy, A. & Hasty, A. H. Weight cycling increases T-cell accumulation in adipose tissue and impairs systemic glucose tolerance. Diabetes 62, 3180–3188 (2013).

21. Cottam, M., Caslin, H., Winn, N. & Hasty, A. Multiomics reveals persistence of obesity-associated immune cell phenotypes in adipose tissue during weight loss and subsequent weight regain. Nature Communications vol. 13 2950 (2022).

22. Netea, M. G., Quintin, J. & van der Meer, J. W. M. Trained Immunity: A Memory for Innate Host Defense. Cell Host & Microbe 9, 355–361 (2011).

23. Kleinnijenhuis, J. et al. Bacille Calmette-Guérin induces NOD2-dependent nonspecific protection from reinfection via epigenetic reprogramming of monocytes. Proc Natl Acad Sci U S A 109, 17537–17542 (2012).

24. Quintin, J. et al. Candida albicans Infection Affords Protection against Reinfection via Functional Reprogramming of Monocytes. Cell Host & Microbe 12, 223–232 (2012).

25. Rodriguez, R. M., Suarez-Alvarez, B. & Lopez-Larrea, C. Therapeutic Epigenetic Reprogramming of Trained Immunity in Myeloid Cells. Trends in Immunology 40, 66–80 (2019).

26. Mulder, W. J. M., Ochando, J., Joosten, L. A. B., Fayad, Z. A. & Netea, M. G. Therapeutic targeting of trained immunity. Nature Reviews Drug Discovery (2019) doi:10.1038/s41573-019-0025-4.

27. van der Heijden, C. D. C. C. et al. Aldosterone induces trained immunity: the role of fatty acid synthesis. Cardiovasc Res 116, 317–328 (2020).

28. Netea, M. G. et al. Defining trained immunity and its role in health and disease. Nat Rev Immunol 20, 375–388 (2020).

29. Kashima, D. T. & Grueter, B. A. Toll-like receptor 4 deficiency alters nucleus accumbens synaptic physiology and drug reward behavior. Proc Natl Acad Sci U S A 114, 8865–8870 (2017).

30. Hao, Y. et al. Integrated analysis of multimodal single-cell data. Cell 184, 3573–3587.e29 (2021).

31. Ge, S. X., Jung, D. & Yao, R. ShinyGO: a graphical gene-set enrichment tool for animals and plants. Bioinformatics 36, 2628–2629 (2020).

32. Orr, J. S., Kennedy, A. J. & Hasty, A. H. Isolation of Adipose Tissue Immune Cells. JoVE (Journal of Visualized Experiments) e50707 (2013) doi:10.3791/50707.

33. Cao, Y. et al. TIDB: a comprehensive database of trained immunity. Database (Oxford) 2021, (2021).

34. Kong, L. et al. Single-cell transcriptomic profiles reveal changes associated with BCG-induced trained immunity and protective effects in circulating monocytes. Cell Reports 37, 110028 (2021).

35. Zhang, B. et al. Single-cell RNA sequencing reveals induction of distinct trained immunity programs in human monocytes. Journal of Clinical Investigation (2022) doi:10.1172/JCI147719.

36. Korbecki, J. & Bajdak-Rusinek, K. The effect of palmitic acid on inflammatory response in macrophages: an overview of molecular mechanisms. Inflamm. Res. 68, 915–932 (2019).

37. Coats, B. R. et al. Metabolically Activated Adipose Tissue Macrophages Perform Detrimental and Beneficial Functions during Diet-Induced Obesity. Cell Reports 20, 3149–3161 (2017).

38. Kratz, M. et al. Metabolic dysfunction drives a mechanistically distinct pro-inflammatory phenotype in adipose tissue macrophages. Cell Metab 20, 614–625 (2014).

39. Seufert, A. L. et al. Dietary palmitic acid induces innate immune memory via ceramide production that enhances severity of acute septic shock and clearance of infection | bioRxiv. (2022) doi:10.1101/2021.06.15.448579.

40. Wager, C. M. L. et al. IFN-γ immune priming of macrophages in vivo induces prolonged STAT1 binding and protection against Cryptococcus neoformans. PLOS Pathogens 14, e1007358 (2018).

41. Edgar, L. et al. Hyperglycemia Induces Trained Immunity in Macrophages and Their Precursors and Promotes Atherosclerosis. Circulation 144, 961–982 (2021).

42. Thiem, K., Stienstra, R., Riksen, N. P. & Keating, S. T. Trained immunity and diabetic vascular disease. Clinical Science 133, 195–203 (2019).

43. Sohrabi, Y. et al. LXR Activation Induces a Proinflammatory Trained Innate Immunity-Phenotype in Human Monocytes. Frontiers in Immunology 11, (2020).

44. Boutens, L. et al. Unique metabolic activation of adipose tissue macrophages in obesity promotes inflammatory responses. Diabetologia 61, 942–953 (2018).

45. Serbulea, V. et al. Macrophage phenotype and bioenergetics are controlled by oxidized phospholipids identified in lean and obese adipose tissue. Proceedings of the National Academy of Sciences 115, E6254–E6263 (2018).

46. Thiem, K. et al. Hyperglycemic Memory of Innate Immune Cells Promotes In Vitro Proinflammatory Responses of Human Monocytes and Murine Macrophages. J.I. 206, 807–813 (2021).

47. Bekkering, S. et al. Oxidized low-density lipoprotein induces long-term proinflammatory cytokine production and foam cell formation via epigenetic reprogramming of monocytes. Arterioscler. Thromb. Vasc. Biol. 34, 1731–1738 (2014).

48. Christ, A. et al. Western Diet Triggers NLRP3-Dependent Innate Immune Reprogramming. Cell 172, 162–175.e14 (2018).

49. Zamarron, B. F. et al. Macrophage Proliferation Sustains Adipose Tissue Inflammation in Formerly Obese Mice. Diabetes db160500 (2016) doi:10.2337/db16-0500.

50. Garcia-Valtanen, P., Guzman-Genuino, R. M., Williams, D. L., Hayball, J. D. & Diener, K. R. Evaluation of trained immunity by β-1, 3-glucan on murine monocytes in vitro and duration of response in vivo. Immunology and Cell Biology 95, 601–610 (2017).

51. Fensterheim, B. A. et al. The TLR4 Agonist Monophosphoryl Lipid A Drives Broad Resistance to Infection via Dynamic Reprogramming of Macrophage Metabolism. The Journal of Immunology 200, 3777–3789 (2018).

52. Owen, A. M., Fults, J. B., Patil, N. K., Hernandez, A. & Bohannon, J. K. TLR Agonists as Mediators of Trained Immunity: Mechanistic Insight and Immunotherapeutic Potential to Combat Infection. Frontiers in Immunology 11, (2020).

53. Kosteli, A. et al. Weight loss and lipolysis promote a dynamic immune response in murine adipose tissue. J Clin Invest 120, 3466–3479 (2010).

