## Supplementary Figures for "Weight cycling induces innate immune memory in adipose tissue macrophages"

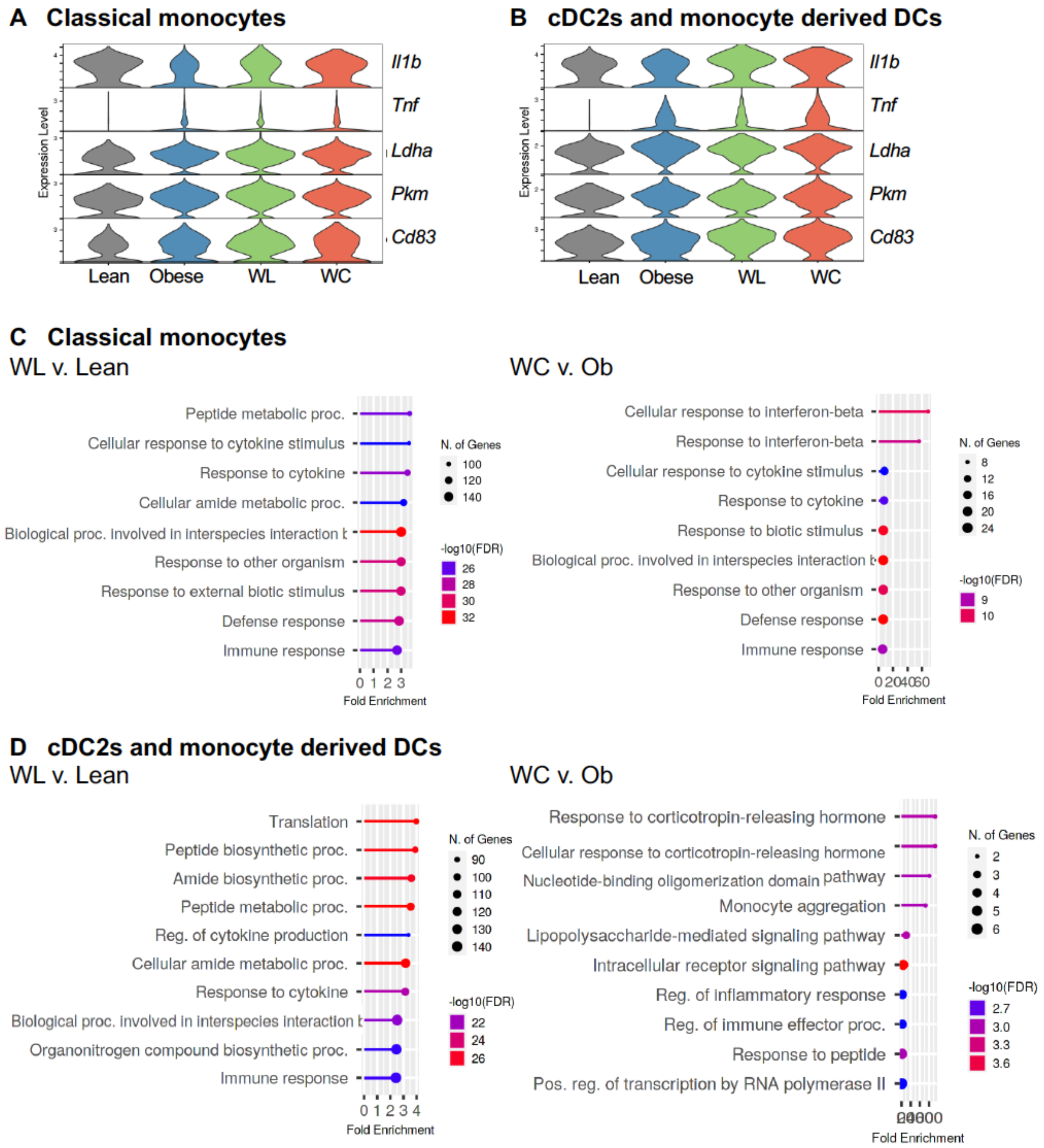


**Supplemental Figure 1: Transcriptional changes in DCs and monocytes.** A) Violin plot of inflammatory (*Tnf, Il1b, Cd83*) and metabolic genes (*Ldha, Pkm*) from lean, obese, weight loss (WL), and weight cycled (WC) in classical monocytes. B) Violin plot of genes by group as in A) for the cDC1 and monocyte- derived DC clusters. C) Pathway analysis of differentially expressed genes from WC vs. obese or WL vs. lean in classical monocytes. D) Pathway analysis as in C) for the cDC1 and monocyte- derived DC clusters.

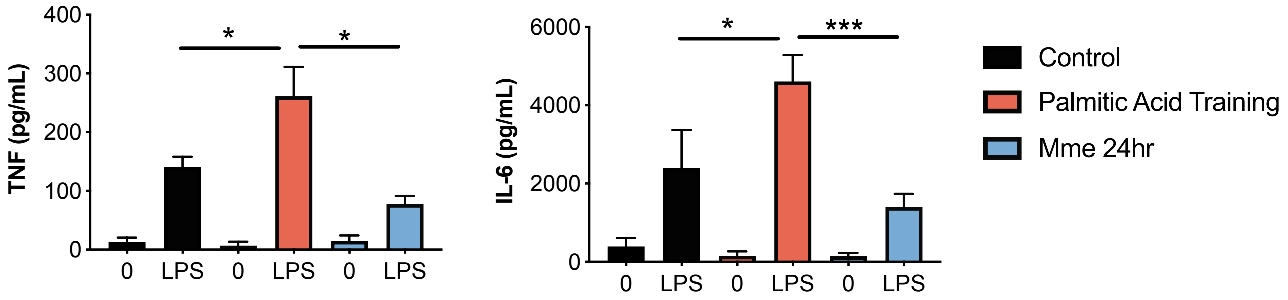


**Supplemental Figure 2: The palmitic acid- induced memory response is different than metabolic activation (MMe).** Bone marrow derived macrophages were treated as in **Figure 2A** with 0.4 mM palmitic acid or treated for 1 day with palmitic acid, glucose, and insulin to induce metabolically activated macrophages (Mme). 100 ng/mL LPS- induced TNF and IL-6 production by ELISA and normalized by protein concentration. Data are means ± SEM of 4 populations, representative of 2 independent experiments.

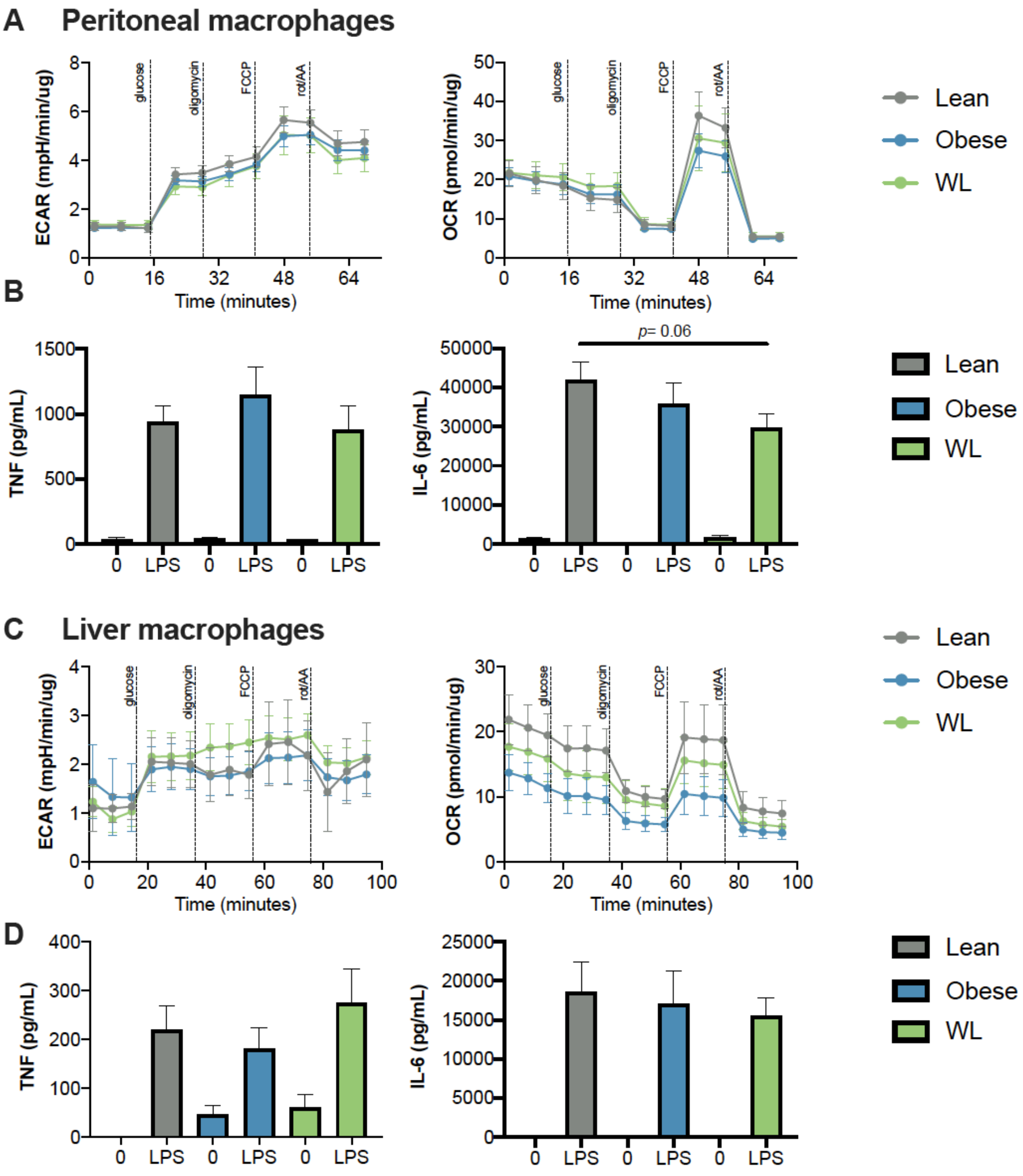


**Supplemental Figure 3: Adipose macrophage inflammation is tissue specific following weight loss.** (A) Extracellular acidification rate (ECAR) and oxygen consumption rate (OCR) during modified mitochondrial stress test by Seahorse metabolic analyzer in lean, obese, and weight loss (WL) peritoneal macrophages selected by 2 hr adherence. **(B)** 100 ng/mL LPS- induced TNF and IL-6 production over 24 hours by ELISA in peritoneal macrophages and normalized by protein concentration. **(C)** Extracellular acidification rate (ECAR) and oxygen consumption rate (OCR) during modified mitochondrial stress test of lean, obese, and WL liver macrophages selected by 2 hr adherence. **(D)** LPS- induced TNF and IL-6 production over 24 hours by ELISA in liver macrophages and normalized by protein concentration. Data are means ± SEM of 6-8 mice.

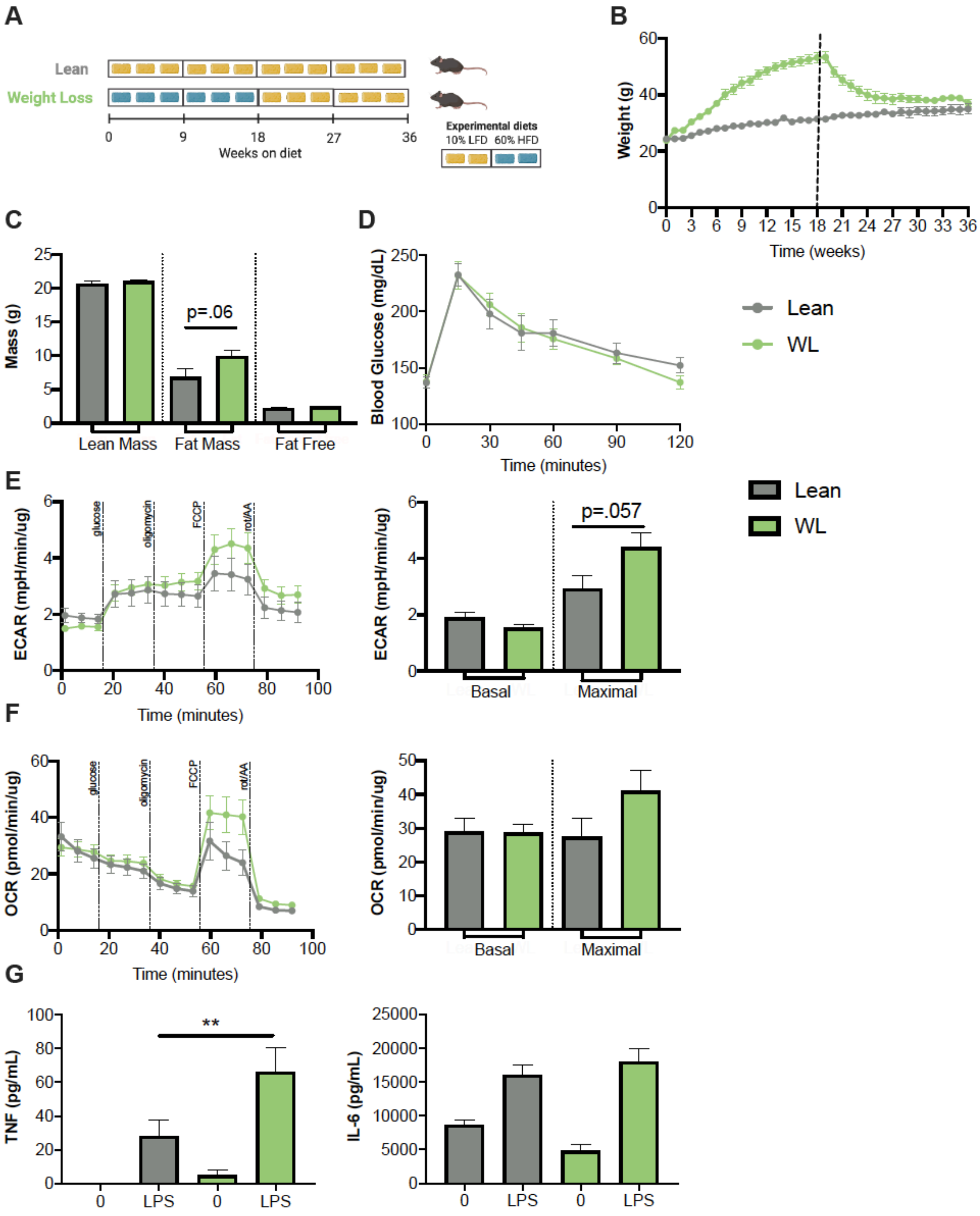


**Supplemental Figure 4: Longer term weight loss (18 wks) does not reverse obesity-associated TNF production. (A)** 27 week weight loss schematic. **(B)** Body mass over time measured weekly with diet switch indicated by dashed lines. **(C)** Lean and fat mass measured by nuclear magnetic resonance. **(D)** Blood glucose during an intraperitoneal glucose tolerance test (1.5 g dextrose/kg lean mass) at 36 weeks. **(E)** Extracellular acidification rate (ECAR) and **(F)** oxygen consumption rate (OCR) of epididymal adipose macrophages selected by 2 hr adherence during modified mitochondrial stress test by

Seahorse metabolic analyzer. **(G)** 100 ng/mL LPS- induced TNF and IL-6 production over 24 hours by ELISA in epididymal adipose macrophages selected by 2 hr adherence and normalized by protein concentration. Data are means ± SEM of 8 mice. **$p$ < .01.

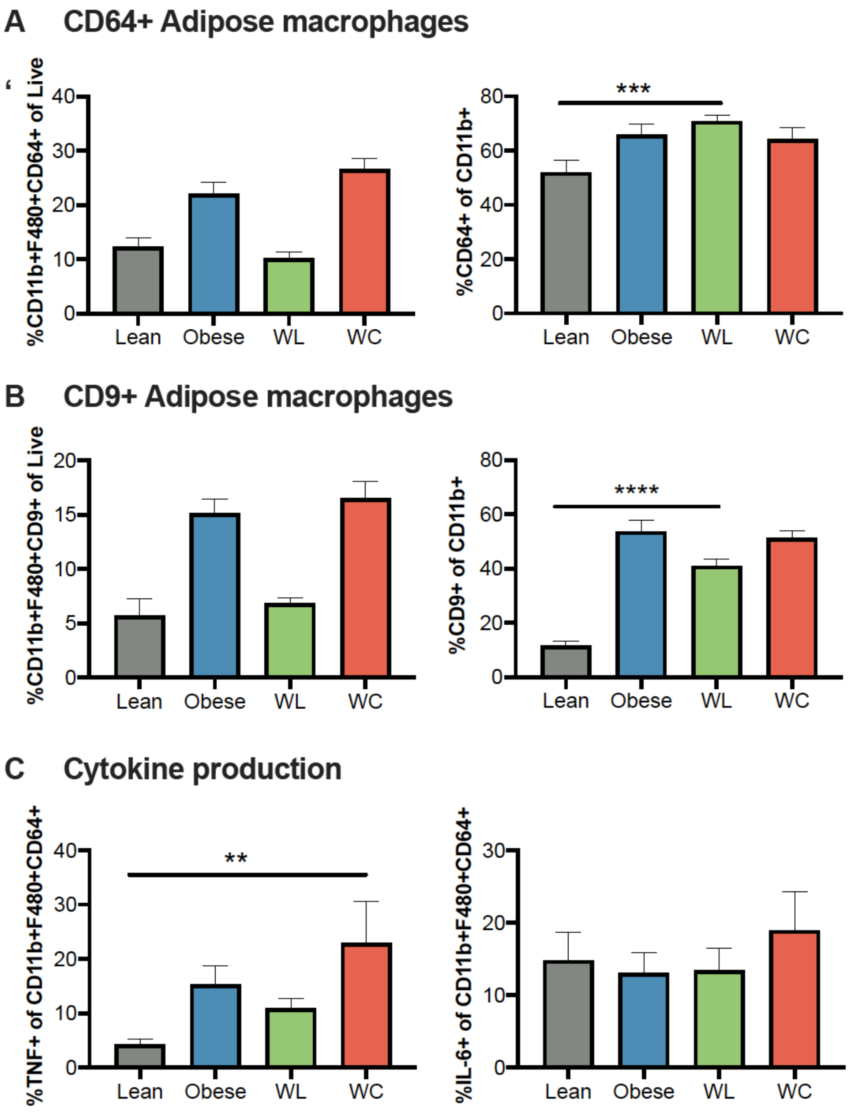


**Supplemental Figure 5: Adipose macrophage populations don't change by flow cytometry with weight cycling despite increased TNF production.** Lean, obese, weight loss (WL) and weight cycled (WC) mice were generated over 27 weeks as in **Figure 1A**. **(A)** CD64+ adipose tissue macrophages (CD11b+, F480+, CD64+) as a percentage of live cells or CD11b+ cells. **(B)** CD9+ adipose tissue macrophages (CD11b+ F480+ CD9+) as a percentage of live cells or CD11b+ cells. **(C)** %TNF+ and %IL-6+ CD64+ adipose tissue macrophages. Data are means ± SEM of 8-12 mice, representative of 2 independent experiments. *$p$ <0.05, **$p$ < .01, *** $p$ <0.001.

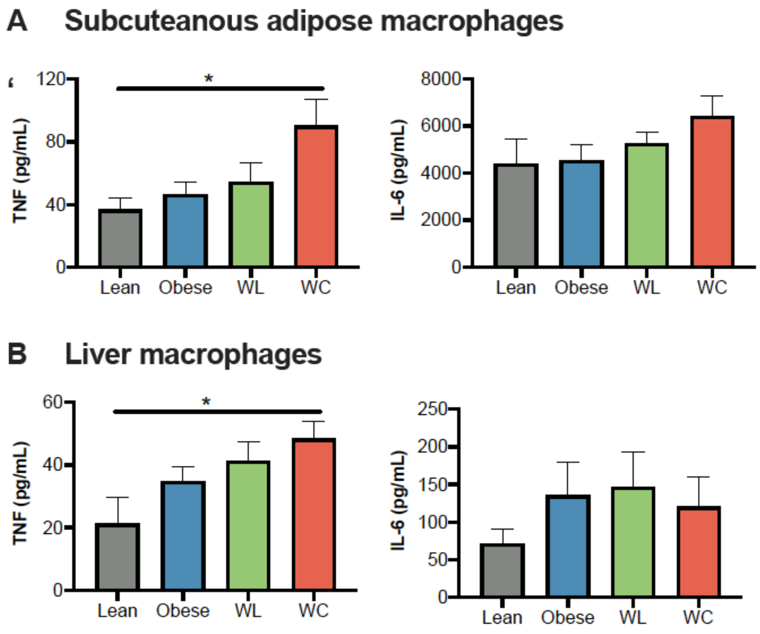


**Supplemental Figure 6: Subcutaneous and liver macrophages also produce more TNF following weight cycling.** Lean, obese, weight loss (WL) and weight cycled (WC) mice were generated over 27 weeks as in **Figure 1A**. **(A)** TNF and IL-6 production over 24 hours by ELISA in subcutaneous adipose macrophages selected by 2 hr adherence. **(B)** TNF and IL-6 production over 24 hours by ELISA in peritoneal macrophages selected by 2 hr adherence. Data are means ± SEM of 6-14 mice, representative of 2 independent experiments. *$p$ <0.05, **$p$ < .01, *** $p$ <0.001
